# Identifying novel age-modulating compounds and quantifying cellular aging using novel computational framework for evaluating transcriptional age

**DOI:** 10.1101/2023.07.03.547539

**Authors:** Chao Zhang, Nathalie Saurat, Daniela Cornacchia, Sun Young Chung, Trisha Sikder, Andrew Minotti, Lorenz Studer, Doron Betel

## Abstract

The differentiation of human pluripotent stem cells (hPSCs) provides access to most cell types and tissues. However, hPSC-derived lineages capture a fetal-stage of development and methods to accelerate progression to an aged identity are limited. Understanding the factors driving cellular age and rejuvenation is also essential for efforts aimed at extending human life and health span. A prerequisite for such studies is the development of methods to score cellular age and simple readouts to assess the relative impact of various age modifying strategies.

Here we established a transcriptional score (RNAge) in young versus old primary fibroblasts, frontal cortex and substantia nigra tissue. We validated the score in independent RNA-seq datasets and demonstrated a strong cell and tissue specificity. In fibroblasts we observed a reset of RNAge during iPSC reprogramming while direct reprogramming of aged fibroblasts to induced neurons (iN) resulted in the maintenance of both a neuronal and a fibroblast aging signature. Increased RNAge in hPSC-derived neurons was confirmed for several age-inducing strategies such as SATB1 loss, progerin expression or chemical induction of senescence (SLO). Using RNAge as a probe set, we next performed an in-silico screen using the LINCS L1000 dataset. We identified and validated several novel age-inducing and rejuvenating compounds, and we observed that RNAage captures age-related changes associated with distinct cellular hallmarks of age. Our study presents a simple tool to score age manipulations and identifies compounds that greatly expand the toolset of age-modifying strategies in hPSC derived lineages.

## Introduction

Aging increases the risk of disease across tissue types. This raises the question of whether targeting the aging process can extend health span and lower the overall disease burden in older adults. Manipulating age is also critical for establishing late-onset disease models using human pluripotent stem cells (hPSC). While hPSC technology offers access to a diverse range of cell types, hPSC-derived lineages represent fetal rather than adult or aged stages and therefore, may not be appropriate for capturing age-dependent disorders without a means to fast-forward cellular age. To develop age-modifying interventions, it is critical to establish methods to quantitatively measure biological age at the cellular level. This is particularly challenging given the complexity of the aging process and its associated biomarkers, which span broad changes at the molecular, cellular, and systemic level.

One major effort in the aging field has been the identification of cellular hallmarks of age^1,2^. Key aging hallmarks include measures of DNA damage, loss of heterochromatin, telomere shortening, loss of proteostasis, mitochondrial dysfunction and induction of senescence among others. Those readouts can be used to demonstrate age-related differences between primary cells derived from either old versus young individuals or to demonstrate the reset of age-related features upon iPSC reprogramming^3,4^. More recently, omics-based clocks have been established to measure cellular age at the molecular level, predict biological or chronological age or to test for changes in rate of aging in an individual^5^. Currently, DNA methylation-based aging clocks, which are based on 5mC methylation status at tissue- and age-related CpG sites^6,7^, dominate the field. Several alternative clocks have been proposed that capture other modalities such as age-dependent changes in transcriptomic, metabolomic or proteomic markers^5^. Transcriptomics-based aging clocks offer the obvious advantage of being widely applicable, easily accessible, and relatively low cost and may provide direct mechanistic insights into the aging process. There have been several efforts to develop both global and cell type specific transcriptional aging clocks^8-11^. To date, the use of these aging clocks has been limited to estimating absolute or relative cellular age. An exciting future application is their use as a direct readout in functional screens aimed at age manipulation to establish improved hPSC-based models of late onset human disease.

Here, we set out to generate a human-centric transcriptomic measure of age that can be used to identify novel aging regulators and to compare mechanisms of aging across different cell types. By performing RNA-seq analysis of matched young versus old human primary fibroblasts, and young versus old brain tissue including frontal cortex and substantia nigra, we identified both conserved and tissue-specific signatures of age. We next developed RNAge, a novel transcriptomic-based method to routinely score changes in the relative age of human cells. We validated our aging score across several independent datasets and showed that it can predict relative age in both cellular rejuvenation and in cellular age induction paradigms. Our scoring approach offers novel insights into the maintenance versus reset of neuronal and fibroblast-specific age signatures during direct and iPSC-based cellular reprogramming. Finally, we used the aging score to query the LINCS L1000 database of cellular perturbations^12^ https://lincsproject.org and to perform an in-silico drug screen for predicted regulators of age in fibroblasts and neurons. Remarkably, this approach identified several novel regulators of age in neurons and in fibroblast that we successfully validated experimentally. Finally, we observed that transcriptional age induction using the various compounds can trigger distinct subsets of the cellular hallmarks of aging. Therefore, RNAge presents a simple strategy to capture distinct cellular mechanisms and temporal dynamics of cellular aging.

## Results

### Identifying transcriptional signatures of age in primary tissues

To identify conserved and tissue specific transcriptional signatures of cellular age we performed bulk RNA sequencing of primary human tissue (fibroblasts, cortex, substantia nigra). A total of 9 t o 10 young (7-14 years old) and 9 old (70-96 years old) samples were sequenced from each tissue type (Figure 1A; Table 1). Samples primarily clustered according to tissue type (Supp. Figure 1A). However, tissue specific clustering of the samples showed a clear separation between young and old samples (Figure 1B), and differential expression analysis identified tissue-specific age-related genes (Figure 1C). Functional enrichment analysis indicated that there is only a very limited functional overlap between age-related gene sets across the tissues (Supp. Figure 1B). In fibroblasts, aging is associated with a downregulation of genes associated with DNA replication, telomere maintenance and cell division and an increase in the expression of genes associated with lipid oxidation amongst others (Figure 1D). These changes are consistent with well described cellular hallmarks of age in proliferating cells such as telomere shortening and cellular senescence leading to decreased cell division and an increase in oxidative stress^1,2^. In contrast, aging in the brain was primarily linked to functional categories associated with neuronal properties and energy and metabolic state. In the cortex, increased age was linked to a decrease in synaptic transmission and plasticity and defects in glutamate receptor signaling. In the substantia nigra, aging was associated with a decrease in neurotransmitter secretion and dopamine transport as well as with changes in cellular respiration, oxidative phosphorylation, and mitochondrial translation (Figure 1D).

**Figure 1:**
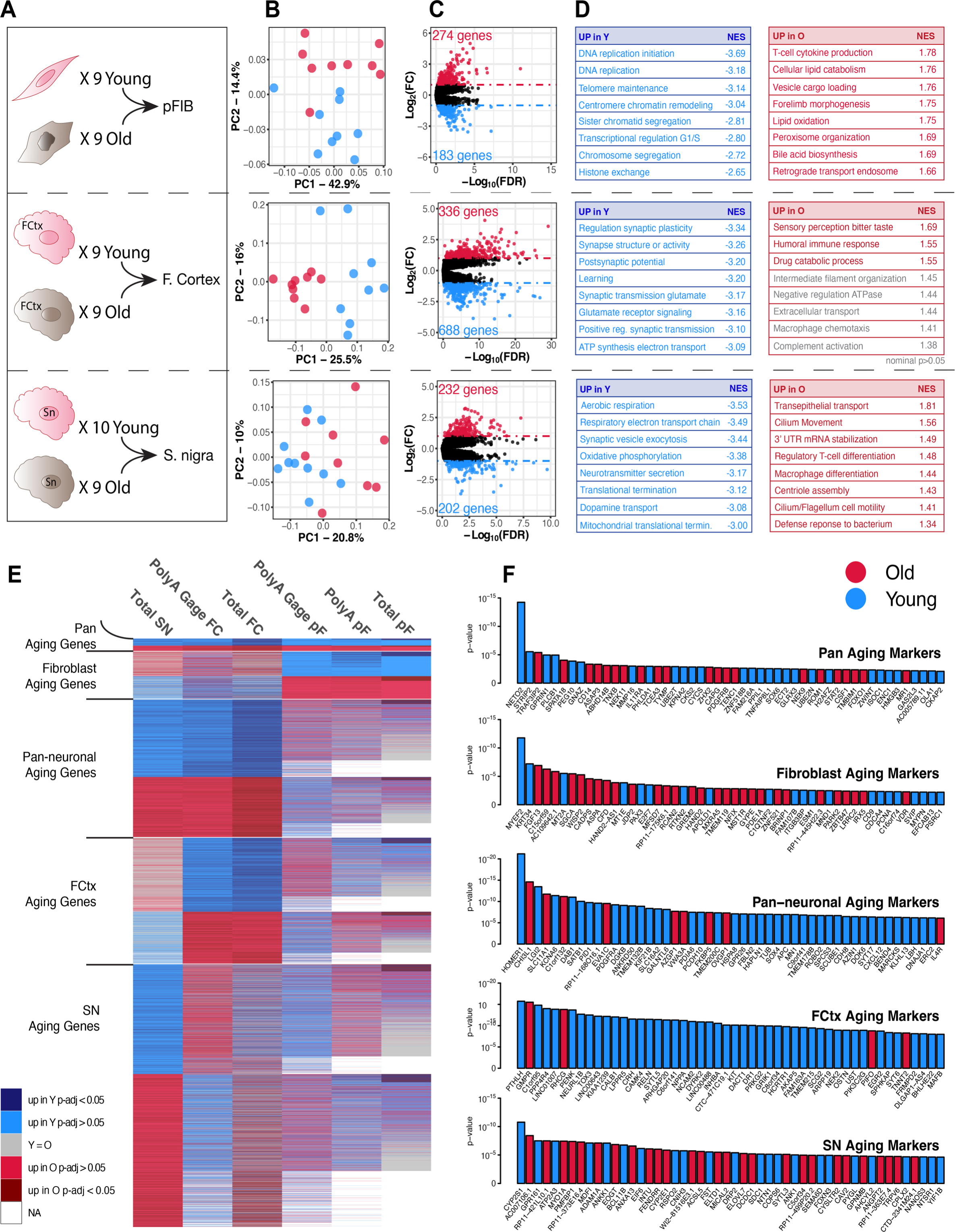
Identification of transcriptional signatures of age in primary fibroblasts, frontal cortex and substantia nigra. **(A)** Schematic outlining of transcriptional profiles of primary cell lines (fibroblasts; pFIB) and tissues (frontal cortex; F. Cortex and substantia nigra; S. nigra) from young (blue) and old (red) donors sequenced in this study. **(B)** PCA of RNA-seq data from primary fibroblasts, frontal cortex and substantia nigra from young (blue) and old (red) donors. **(C)** Volcano plots showing significantly differentially expressed genes (FDR <0.1) between young and old primary tissues. Genes in blue are significantly higher expressed in tissues from young donors and in red are significantly higher expressed in tissues from old donors. **(D)** Functionally enriched pathways of age-associated genes in each tissue (selected terms) and their respective Normalized Enrichment Score (NES). **(E)** Heatmap indicating directionality of the change in gene expression in young vs old primary tissue samples. Genes in blue are more highly expressed in young and those in red are more highly expressed in old. Darker shades indicate that change is statistically significant (up to FDR <0.05). Genes whose expression is unchanged are indicated in grey and those not expressed in a specific tissue are in white. RNA-seq datasets generated as part of this study were: Total SN (substantia nigra), Total FC (frontal cortex), Total pF (total RNA of primary fibroblast) and PolyA pF (poly-A selection of primary fibroblast). External datasets were PolyA Gage FC (poly-A selection of frontal cortex) and PolyA Gage pF (polyA selection of primary fibroblast). **(F)** Top 50 genes for each subcategory of aging genes ranked according to combined p-value. Genes in blue are upregulated in young tissues and in red are upregulated in old tissues.

Our aim is to utilize these age-related, dysregulated gene expression patterns to develop a scoring model that can be used to assess rejuvenation or induction of cellular age. To this end, we incorporated external datasets^13^, as well as internal poly-A RNA-seq data (Table 1) from a subset of our fibroblast samples. To overcome the large technical variability introduced by different experimental platforms and independent studies^5,9,11^ we identified the subset of genes whose expression change in a consistent direction across the multiple datasets. We identified a set of five mutually exclusive aging signatures. These signatures include pan-aging genes that mark aging across all three lineages, fibroblast specific, pan neuronal, frontal cortex (FC) specific and substantia nigra (SN) specific (Figure 1E). For all categories, the top 50 genes (ranked by combined p-value) are presented in Figure 1F with genes that are decreased in old versus young samples indicated in blue and those increased in old versus young indicated in red. We also performed pathway enrichment analysis for the genes in each of the different signatures to identify the cellular pathways that might underpin each aging signature (Supp. Figure 1C-G). The pan-aging gene set was associated with an increase in innate immunity in aged samples and decreased protein folding, protein stabilization and processing of DNA double stranded breaks. The fibroblast aging signature was largely associated with a decrease in pathways associated with cell division and cell cycle, consistent with the functional enrichment in Figure 1D. Pan neuronal genes that were downregulated with age were linked to learning, synapse assembly and transmission whereas genes upregulated with age were associated with triglyceride catabolism as well as pathways related to cellular stress including: response to lipopolysaccharide, cellular response to hydrogen peroxide and cellular response to glucose starvation. Interestingly, in the fibroblast specific, FC specific and SN specific signatures there was an increase in functions linked to differentiation or development in aged tissues. This is compatible with previous studies suggesting that age is associated with a loss of cellular identity^14 15^.

### Establishment and validation of RNAseq-based aging scores

Using the six age-associated datasets we established a computational pipeline (“RNAge”) to measure changes in relative age between two groups of samples. For example, upon a genetic perturbation in fibroblasts, one can assess whether the resulting gene expression changes are consistent with the changes measured between fibroblasts from young and old donors. To establish our score, we used the aging gene signature panels (Pan, Fibroblast, Pan Neuronal, FC and SN) to identify a subset of age-regulated genes that are either upregulated or downregulated in young vs. aged cells. For each subcategory, we include the top 100 genes ranked according to their combined p-value in the corresponding tissues (Figure 1E). Briefly, we first calculate the t-score of the age signature genes between contrasting two conditions in the test set. The t-scores are then multiplied by a binary vector indicating the expected expression change for each gene (i.e. up in old or young) and the resulting weighted t-score values are averaged for the final RNAge score. (Figure 2A, see method section). We also calculate the percent of genes of the age signature that change expression in the expected direction to assess if the aging score is driven by very large changes in only a small subset of the age associated genes or by a broader change across most of the genes.

**Figure 2:**
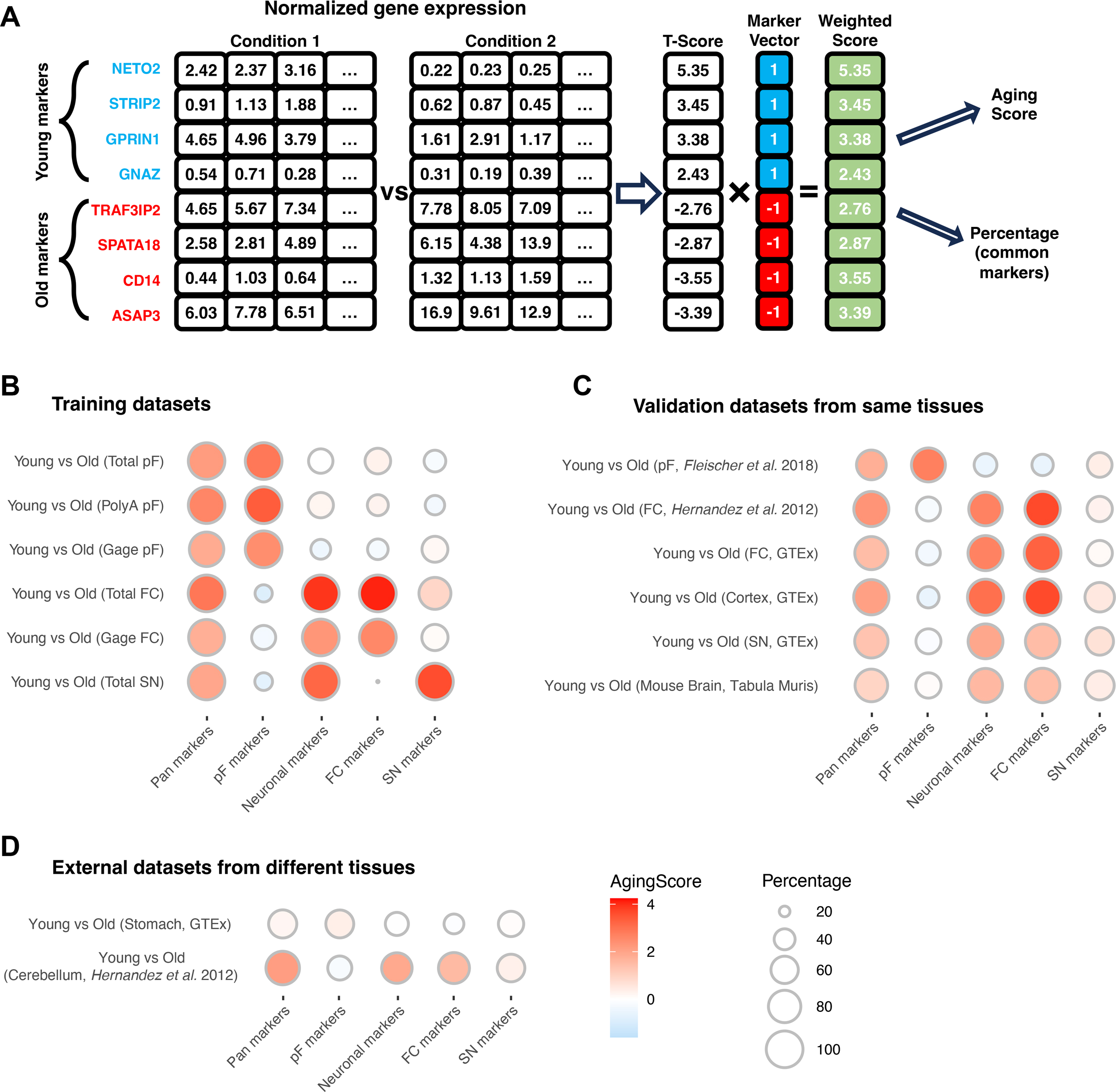
Establishment and validation of RNAge score. **(A)** Schema outlining the steps to calculate the RNAge score. Welch’s two-sample t tests are calculated on the normalized values of the top 100 age signature genes (Figure 1E) between two conditions. The t-test scores are weighted by multiplying a binary marker vector that signifies the directionality of the expected expression changes (i.e., negative for genes marking older samples and positive for genes expressed highly in younger samples) and the mean values is the final age score. Percentage of common genes is the fraction of genes with positive weighted t-score. **(B)** RNAge scores of the datasets used to establish aging signature subcategories. RNA-seq data used in Total pF, PolyA pF, Total FC and Total SN were generated as part of this study. Gage pF and Gage FC are similar young and old samples that were profiled independently in a different study^13^. RNAge score refers to the modified t-score for each pairwise comparison and is indicated by the color of the bubbles. The size of the bubbles indicates the percentage of genes that make up the aging score that change in the expected direction for young vs old. **(C)** Independent validation of the RNAge score for pF, FC, Cortex, SN, and mouse brain from additional external studies. Tissue type, species and data source are indicated on the bubble plot. **(D)** Application of RNAge score to young and old human primary stomach and human primary cerebellum from previously published data.

Applying our aging score to the original datasets used to develop the scoring method confirmed the specificity of the scores across all signatures (Figure 2B). To test the utility of the score, we applied the aging score on several independent aging datasets that were not used in the development of RNAge (Figure 2C). Our aging score performed well in assessing independent transcriptomic datasets from old versus young human primary fibroblasts^9^ and in three independent datasets of young versus old human frontal cortex^16,17^. The Hernandez et al. dataset was generated using microarray-based technology indicating that our aging score is applicable across different gene expression platforms. Finally, our aging score was also capable of measuring changes in the relative age of young versus old mouse brain further highlighting the broad utility and robustness of our score, even across species. The SN-specific age score was less robust than the other scores as indicated by the fact that when we scored age in the GTEx SN dataset we could detect an increase in cellular age in the pan aging, pan neuronal and FC scores but only a marginal increase in the SN specific score (Figure 2C). This is largely a reflection of a weak signature that was derived from only a single dataset in our original SN samples.

Finally, we tested whether our aging score was applicable to other cell types (Figure 2D). In the cerebellum^16^, our pan aging and pan neuronal scores were predictive of cellular age indicating that those scores may be broadly useful across neuronal subtypes. However, none of the scores could discriminate between young versus old primary stomach samples^17^. Those data further illustrate the tissue-specificity of the transcriptional aging scores and raise the question whether including additional samples across different organ systems would lead to a truly universal transcriptional aging score across all cell types of the human body.

### ‘RNAge’ score can be used to measure cellular rejuvenation

Another way to validate our aging score is to test its effectiveness in scoring rejuvenation paradigms. Transcriptional reprogramming of primary fibroblasts to pluripotent stem cells is one of the best described methods of resetting cellular age^3,4,18^. Therefore, we took a subset of our old versus young primary fibroblast cellines that we had used to establish our RNA-based aging score, reprogrammed them back to pluripotency and then differentiated those stem cells back into iPSC-derived fibroblasts (Figure 3A, Supp. Figure 2A). After confirming the successful reprogramming and re-differentiation to fibroblasts both via immunohistochemistry and flow cytometry for canonical fibroblast cell surface markers (Figure 3B), we performed RNAseq. Those gene expression data confirmed the loss of fibroblast identity in iPSC and reestablishment of a fibroblast identify upon re-differentiation (Figure 3C). Next, we tested whether DEGs between young and old primary fibroblasts were maintained during reprogramming and re-differentiation. As expected, reprogramming fibroblasts to iPSCs resulted in the loss of age-related DEGs and differentially methylated genes (DMGs), and those DEGs and DMGs were not re-established once iPSCs were differentiated back into iPSC-derived fibroblasts (Figure 3D and Supp. Figure 2B-E). The loss of age-related transcriptional signatures can be illustrated by focusing on the histone family of genes. These genes are amongst the most significantly differential transcripts between young versus old primary fibroblasts in agreement with several studies showing a global reduction in histone proteins with age^19^. Consistent with this, we found that *HIST1H2BH, HIST1H4F, HIST3H2BB* and *HIST1H3E* expression were reduced in old fibroblasts compared to young fibroblasts and this differential expression is erased in iPSCs and in iPSC-derived fibroblasts (Supp. Figure 2F). After demonstrating that our reprogramming and re-differentiation platform rejuvenates fibroblasts from old donors, we applied the aging score to the iPSC-derived fibroblasts and compared the expression relative to the matched primary fibroblasts they were derived from. Young and old primary fibroblasts show an increased aging score relative to iPSC derived fibroblasts with a more pronounced effect when comparing the old primary fibroblasts to iF (Figure 3E). As expected, there was no difference in the relative age of the iPSC-derived fibroblasts despite the large difference in age of the original donors. Together, these results support previous findings that aging signatures are erased during reprogramming^4,20^ and validates the use of our aging score in measuring cellular rejuvenation.

**Figure 3:**
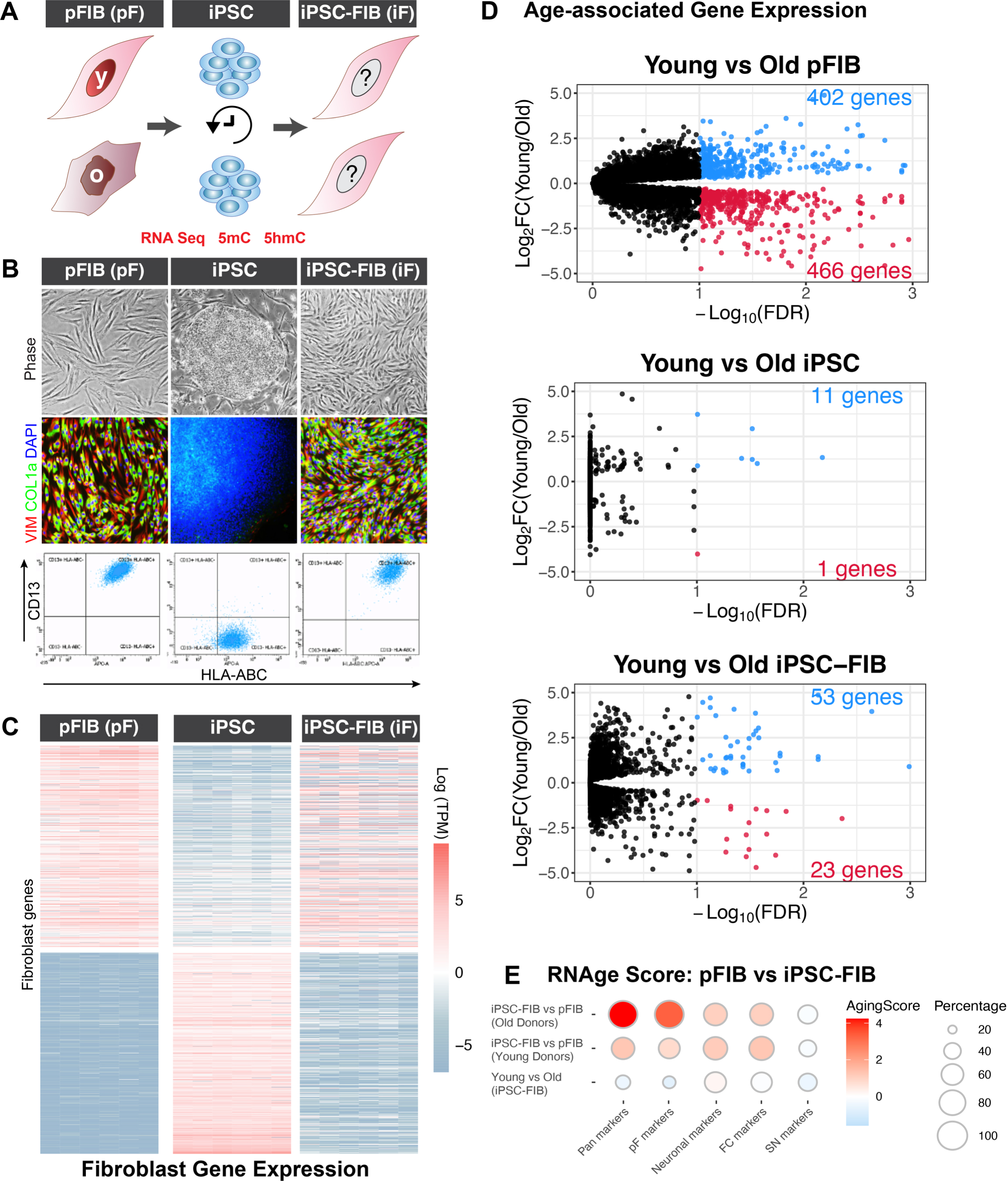
RNAge score can be used to measure cellular rejuvenation. **(A)** Experimental schema of Fibroblast rejuvenation experiment. Primary fibroblasts (pFIB; pF) from young and old donors were first reprogrammed to pluripotent stem cells (iPSC) then re-differentiated into iPSC derived fibroblasts (iPSC-FIB). RNA, 5mC and 5hmC was performed at each step. **(B)** Morphology, immunofluorescence for the fibroblast markers COL1A and VIM and flow cytometry for CD-13 and HLA-ABC in primary fibroblasts, iPSCs generated from these fibroblasts and in matched iPSC-derived fibroblasts. **(C)** RNA-seq analysis showing that gene expression of fibroblast specific genes in iPSC-FIB closely resembles that of pFIB indicating the re-establishment of fibroblast identity in iPSC-FIB. (N=6 independent cell lines at each stage; 3 derived from young donors and 3 derived from old donors). **(D)** Volcano plots showing DEGs (FDR<0.1) between young and old pFibs, iPSCs derived from young and old donors and iPSC-FIB generated from these stem cells. The loss of aged signature (rejuvenation) is indicated by the reduction in the number of significantly differentiated genes in iPSC-FIB compared to pF. (N=3 independent cell lines derived from young donors and N=3 derived from old donors). **(E)** RNAge score comparing the relative age of the iPSC-FIBs (iF) to their matched pFIBs (pF) or direct comparison of the relative age of iFs from young or old donors. The largest RNAge score are when comparing iPSC-FIB to their matched primary fibroblast, whereas RNAge score from young vs. old iPSC-FIB is minimal (N=3 independent cell lines derived from young donors and N=3 derived from old donors)

### RNAge score can successfully identify manipulations that induce cellular age

Progeroid diseases represent widely used genetic models of cellular aging. However, there has been a long-standing debate about whether progeroid diseases phenocopy chronological aging or involve distinct molecular mechanisms and the induction of distinct cellular hallmarks of age^21^. To test this we performed RNA, 5mC and 5hmC profiling on Hutchinson-Gilford progeria syndrome (HGPS) fibroblasts (age 3-14; Table 1) and compared the RNA expression, including RNAge, and DNA methylation profiles of these samples to those of young and old control primary fibroblast samples. PCA analysis indicated that HGPS fibroblasts clustered separately from the old primary fibroblasts This was true for both the transcriptome (Figure 4A) and the methylation profiles (Supp. Figure 3A and B). We also applied our aging score to directly to compare the cellular age of the HGPS fibroblasts to young fibroblasts (Figure 4B). Interestingly, we saw no increase in cellular age in HGPS fibroblasts. This finding was further confirmed in similar data published by a prior independent study of HGPS fibroblasts^9^ (Figure 4B). Together, the transcriptional differences between HGPS and primary fibroblasts and the low aging score between young versus HGPS fibroblasts suggests that HGPS induces a disease-specific aging signature rather than physiological age in fibroblasts.

**Figure 4:**
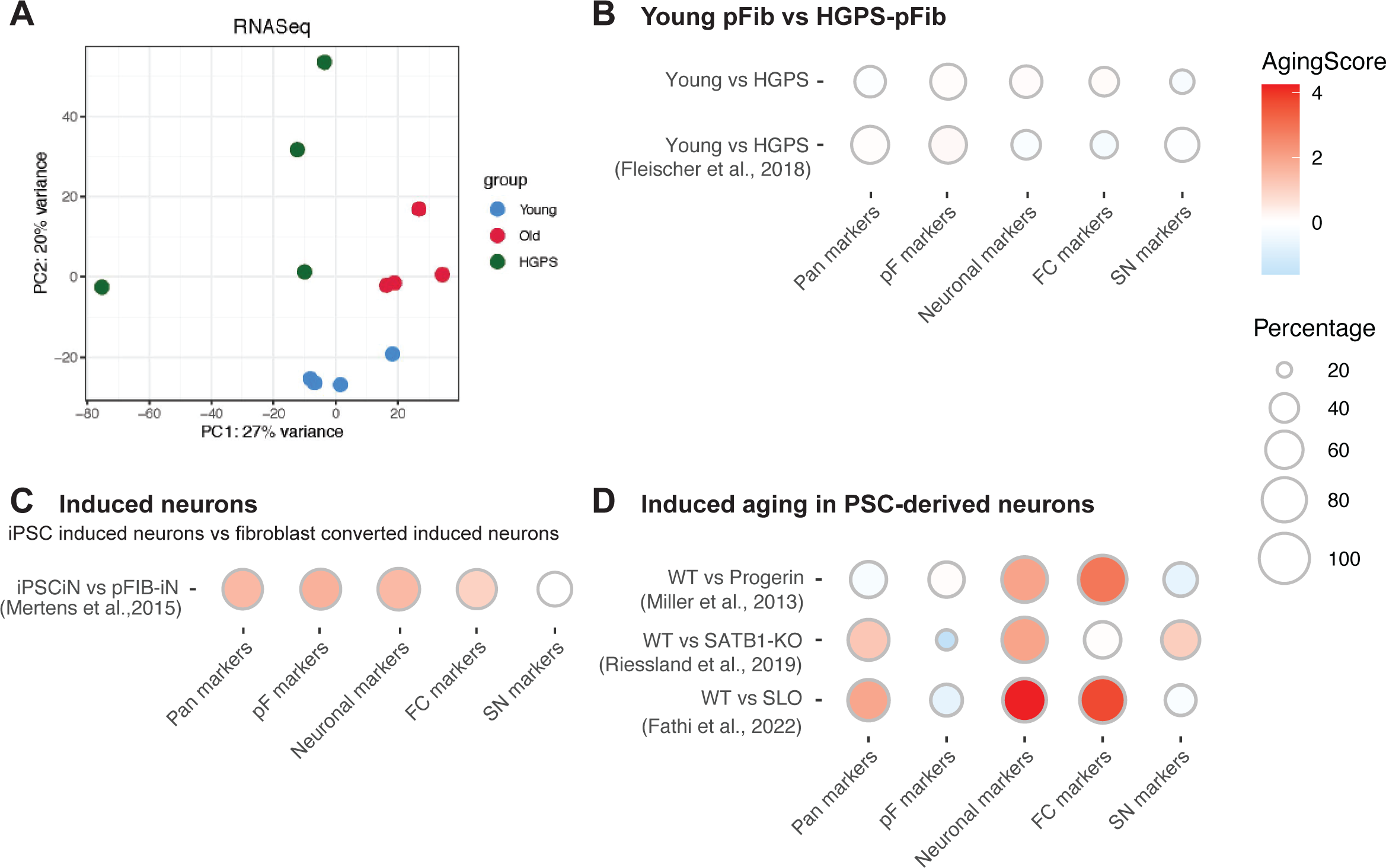
RNAge score in accelerated and induced aging models. **(A)** PCA of RNA-seq data from primary fibroblasts originating from young, old or individuals with HGPS. **(B)** RNAge score of young pFIBs vs. HGPS pFIBs form this study data (top) and similar comparison using RNA-seq data from Fleischer et. al., 2018 (bottom) demonstrating that HGPS do not mimic natural aging. **(C)** RNAge score of iPSC derived induced neurons (iPSCiN) vs. induced neurons derived from primary fibroblasts (pFIB-iN) from Mertens et al., 2015 indicating that trans-differentiated neurons from aged fibroblasts maintain both neuronal and fibroblast aging. **(D)** RNAge measurement in PSC-derived neurons in response to multiple published induced aging strategies; Progerin expression, SATB1-KO, and SLO (Miller et al 2013, Riessland et al 2019 and Fathi et al 2022)

One alternative approach for studying the mechanisms of aging in neurons is to use direct fibroblast conversion to induced neurons (iNs). These neurons are generated by the ectopic expression of key neuronal transcription factors, such as Ngn2 and Ascl1, in primary fibroblasts from individuals of varying age^13,22,23^. When iNs are derived from fibroblasts of old individuals, several studies reported that they retain a transcriptional aging signature and show age-associated disease phenotypes in models of neurodegeneration^13,14,24^. Therefore, we wanted to test whether our score could detect an increase in cellular age in iNs generated from old fibroblasts relative to iNs generated from PSCs where age related signatures are erased during the reprogramming process. Our aging score showed that forebrain neurons generated from old fibroblasts displayed an increase in the pan aging, neuronal aging and FC-specific aging score but not the SN aging score. However, unexpectedly, these fibroblast-derived neurons also maintained their fibroblast-specific aging score in iNs (Figure 4C), a score not present in neurons that have undergone chronological aging (compare to Figure 2B,C).

Another major challenge in aging research is having a reproducible way to score age following various age modifying treatments and to compare the efficiency of age induction across conditions. Such age induction strategies are critical for the study of late onset diseases in hPSC-based models and may open new avenues for identifying and validating novel therapeutic interventions. To date, at least three distinct age induction strategies have been described in neurons including: the ectopic expression of Progerin^4^ (responsible for HGPS), the knockout of SATB1 in dopaminergic neurons resulting in increased cellular senescence^25^ and the chemical induction of senescence in fibroblast and neurons using the “SLO” small molecule cocktail^26^. Using our novel age scoring methods, we observed that the ectopic expression of Progerin in iPSC-derived neurons triggered a pan neuronal and FC aging signature but not an increase in the pan aging score (Figure 4D). This is consistent with our results from primary HGPS fibroblasts that did not show a pan aging, or fibroblast aging score (Figure 4B). In addition to inducing cellular aging hallmarks and phenotypes consistent with neurodegenerative disease as reported in previous studies, we observe that SATB1 KO and SLO treatment induce a robust increase in the transcriptional aging score in each of the expected categories (Figure 4D)

### RNAge score is tractable and can be used to identify age-regulating compounds

Our study demonstrates that RNAge can be used to compare changes in cellular age across primary tissues and in response to different aging and rejuvenating paradigms. We next wanted to test whether it could also be used as a drug discovery tool to identify potential drivers of age. To achieve this goal, we designed an in-silico screen (outlined in Figure 5A) using the LINCS L1000 dataset that represents a large transcriptional dataset following > 1 million pharmacological or genetic perturbations in a diverse set of cell lines^12^. Given the evidence for cell-type specificity of the aging signatures presented, we restricted our screen to a smaller subset of perturbations performed in the cell types that are most similar to the primary tissues used to derive our fibroblast and neuronal age signatures (1HAE and NEU). 1HAE is a normal fibroblast line while NEU corresponds to iPSC derived neurons. We further restricted our analysis to drug-based perturbations as these will be more straightforward to validate and more amenable for widespread use as a tool in iPSC-based disease modeling. By restricting our analysis to those parameters, the L1000 dataset included a total of 348 and 3968 perturbations for 1HAE and NEU respectively. We next calculated the aging scores for each of those perturbations relative to matched control samples in the dataset and averaged them to get a single value for each perturbation (Figure 5B).

**Figure 5:**
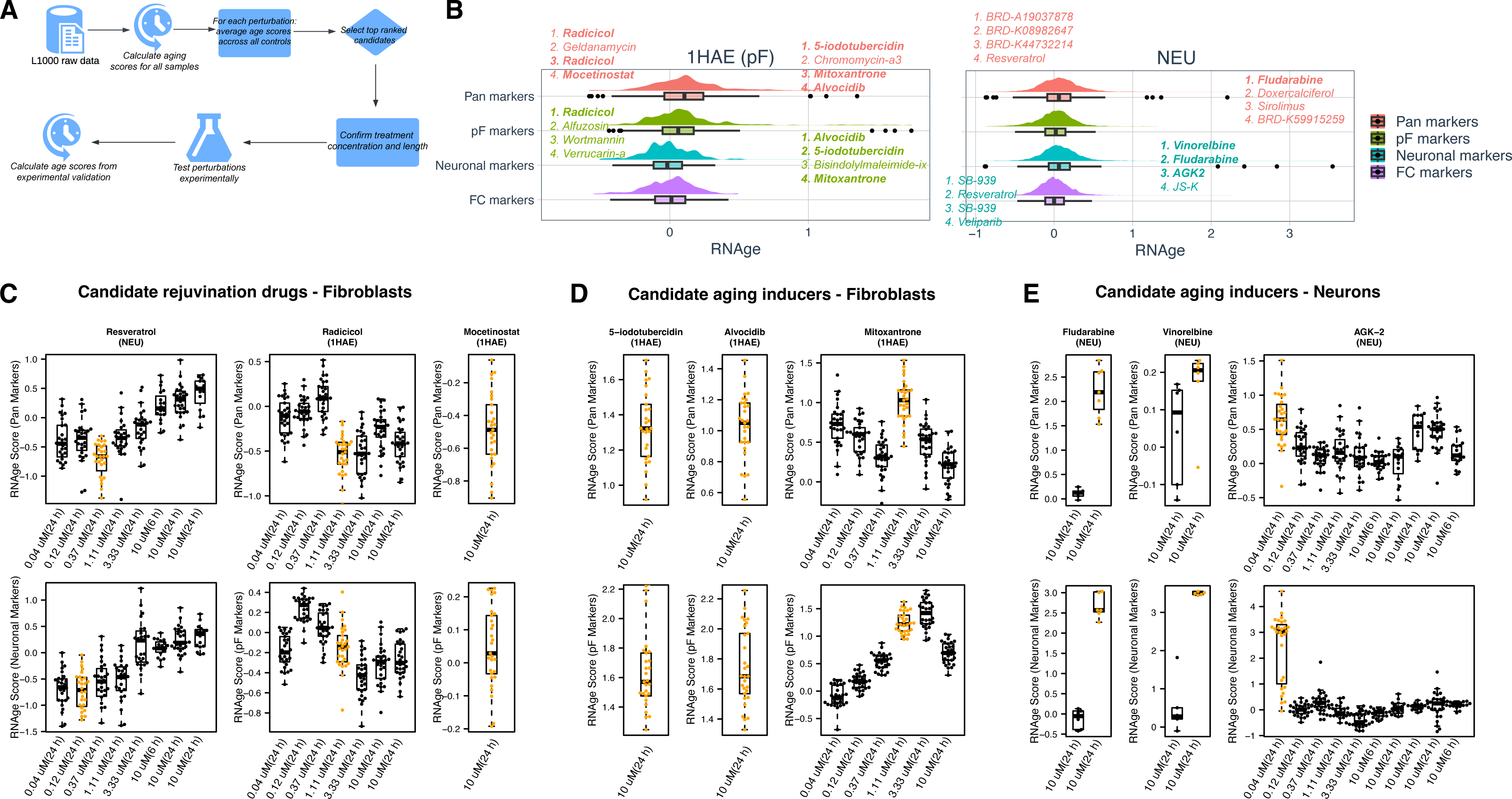
Identification of novel regulators of cellular age using RNAge score. **(A)** Workflow used to score all perturbations in the LINCS L1000 dataset using RNAge. Top scoring perturbations were then tested experimentally. **(B)** Distribution of the RNAge scores for all drug-based perturbations in the L1000 library that were performed in fibroblasts (1HAE; left panel) and neurons (NEU; right panel). The top ranked compounds according to the pan aging and fibroblasts scores are indicated on the fibroblast (1HAE) plot and the top ranked compounds according to the pan aging and pan-neuronal scores are indicated on the neurons (NEU) plot (see points and listed compound names). Perturbations selected for experimental validation are in bold. **(C-E)** Calculation of RNAge for every concentration of our candidate age regulators available within the LINCS 1000 dataset. The RNAge score was calculated relative to every control sample within the dataset. Conditions with the most pronounced effect (yellow) were selected for downstream validation.

Aging scores fell on a normal distribution for all cell types and sub-scores (Figure 5B) indicating that most drug treatments did not induce changes in the aging score. We selected a subset of the top scoring age inducers and rejuvenators for experimental validation. As candidate factors to induce age in fibroblasts we selected Alvocidib (CDK inhibitor), 5-iodotubercin (adenosine kinase inhibitor) and Mitoxantrone (Topoisomerase II inhibitor) for experimental validation, as these three drugs are included in the top four age inducers for both the pan aging and the fibroblast aging scores. We selected Fludarabine (STAT1 and DNA synthesis inhibitor), Vinorelbine (microtubule antagonist) and AGK-2 (SIRT2 inhibitor) for further validation in neurons, as these were the top three candidate regulators of the pan-neuronal score. Fludarabine was also the top candidate as measured by the pan-aging score.

Although, our study focused on age induction strategies due to their potential in aiding the modeling of late onset disease, we also addressed whether our screening strategy can identify candidate rejuvenating drugs. We performed these experiments in fibroblasts due to the availability of primary fibroblasts isolated from aged individuals which is not readily feasible for primary human neurons. We selected Radicicol (HSP90 inhibitor), Mocetinostat (HDAC inhibitor) and Resveratrol (multiple functions) for experimental validation. Radicicol is the top ranked rejuvenation candidate when looking at both the pan and fibroblast aging scores whereas Mocetinostat is the top ranked rejuvenator of the pan aging score and has an independent mechanism of action from Radicicol. We also included resveratrol, a drug that was not included in the LINCs dataset for the 1HAE (fibroblast) line but was a top scoring rejuvenator in the neuron dataset and is widely described as a candidate drug promoting longevity^27,28^.

There are multiple independent perturbations for many of our candidate age regulators within the L1000 dataset. These perturbations represent replicate experiments and different drug concentrations or treatment lengths. To identify the optimal experimental conditions for validation we calculated the aging score for every perturbation associated with our selected candidates relative to every control sample available within the dataset (Figure 5C-E). In some cases, this generated an in-silico concentration curve for candidate age regulators allowing us to select the predicted optimal concentration for each drug. To maintain consistency with the L1000 data generation, cells were harvested following 24hr of the drug treatment. Principal component analysis (Figure 6A) indicated that each of the fibroblast rejuvenation candidates had a strong transcriptional effect relative to their respective vehicle controls (DMSO for Mocetinostat and Resveratrol and Ethanol for Radicicol). Likewise, all the fibroblast aging inducers also segregated from the control, with Alvocidib and 5-iodotubercin clustering closely together. However, the AGK-2 treated neurons clustered with the control sample suggesting that it may not have any obvious effect under the conditions tested. Based on the L1000 data the induction of an aging score in response to AGK-2 treatment is highly sensitive to concentration and it is possible that the optimal concentration or treatment differed in our system (Figure 5E).

**Figure 6:**
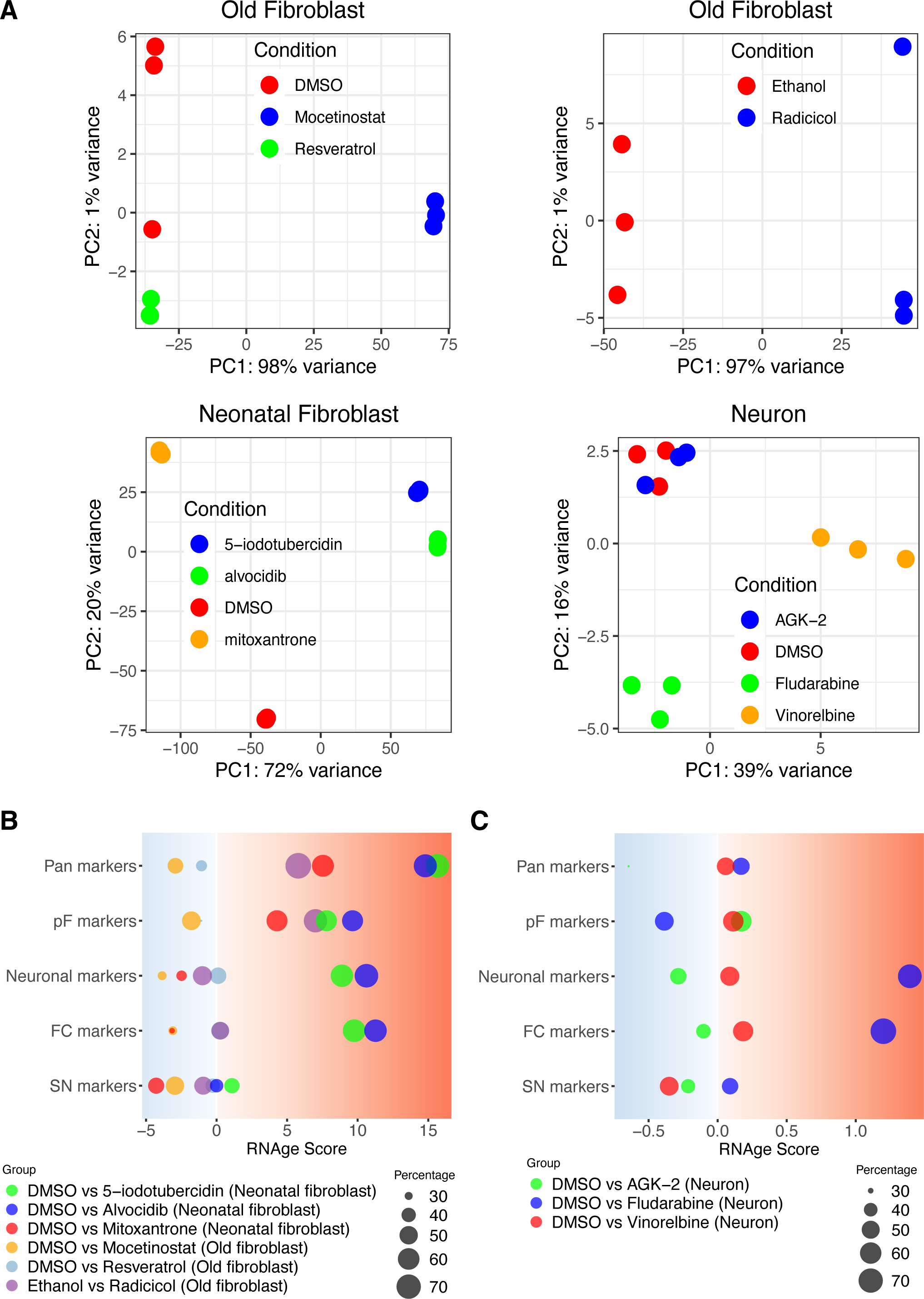
Experimental validation of top scoring modulators of RNAge. **(A)** PCA of RNA-seq data from old fibroblasts treated with candidate rejuvenating compounds and respective controls (top row) and young fibroblasts and neurons treated with candidate age inducers (bottom row). **(B)** RNAge score of old and fetal fibroblast treated with age inducers or rejuvenation agents. Note that the size of the bubble indicates the percent of genes that change in the expected direction between young and old. Therefore, a larger bubble indicates a greater level of agreement with increased age and a smaller bubble shows smaller agreement with age induction and, consequently, a greater agreement with cellular rejuvenation. For example, Resveratrol has a very small bubble size in the fibroblast aging score indicating gene expression changes that are highly correlated with rejuvenation. **(C)** RNAge score of young, iPSC-derived neurons treated with age inducers.

We then used the RNA-seq data to calculate the RNAge score for each of the perturbations relative to their controls (Figure 6B-C). Importantly, Mocetinostat was able to reverse cellular aging in old primary fibroblasts (Figure 6B). Resveratrol showed a similar but more muted effect towards rejuvenation. However, Radicicol did not trigger rejuvenation but rather showed evidence of inducing cellular aging in fibroblasts (pan and fibroblast aging scores). In contrast, treatment of fetal human dermal fibroblasts (HDF-f) with all three of our candidate drugs resulted in an increased age score confirming our initial predictions. Alvocidib and 5-iodotubercin, which clustered together on the PCA plot resulted in a very similar age score induction profile whereas mitoxantrone was distinct and only showed induced aging in the pan aging and fibroblast categories. As predicted by the transcriptional similarities of DMSO treated and AGK-2 treated neurons (Figure 6A), AGK-2 had no impact on aging in neurons under our experimental conditions (Figure 6C) while Fludarabine strongly induced aging in neurons based on both the pan neuronal and FC sub scores as well a weak induction of the pan aging score. Vinorelbine treated neurons showed only slight, albeit consistent, induction of aging scores across the aging signatures.

### Correlation of RNAge to the hallmarks of cellular age

To date, measuring the ‘hallmarks of age’ has been one of the primary ways of quantifying the impacts of putative regulators of aging^1,2^. Here, we developed an independent aging score (RNage) based on transcriptional changes and used it to identify novel regulators of cellular age. Next, we wanted to assess whether the age modifying drug treatments also trigger cellular hallmarks of age and whether those hallmarks are consistent across perturbations and cell types. We developed a robust platform to quantify several hallmarks of age in parallel by immunocytochemistry and high content image analysis (Figure 7A). The experimental aging hallmarks^1,2^ include loss of heterochromatin (H3K9me3 and LAP2), disrupted nuclear lamina (decreased nuclear roundness and loss of LAP2), increased DNA damage (γH2AX), increased cellular senescence (p21 and nuclear area), disrupted proteostasis and increased dependence on autophagy (BAG3^29,30^) and increased CAV1 (multiple functions, broadly increased with age^31-34^).

**Figure 7:**
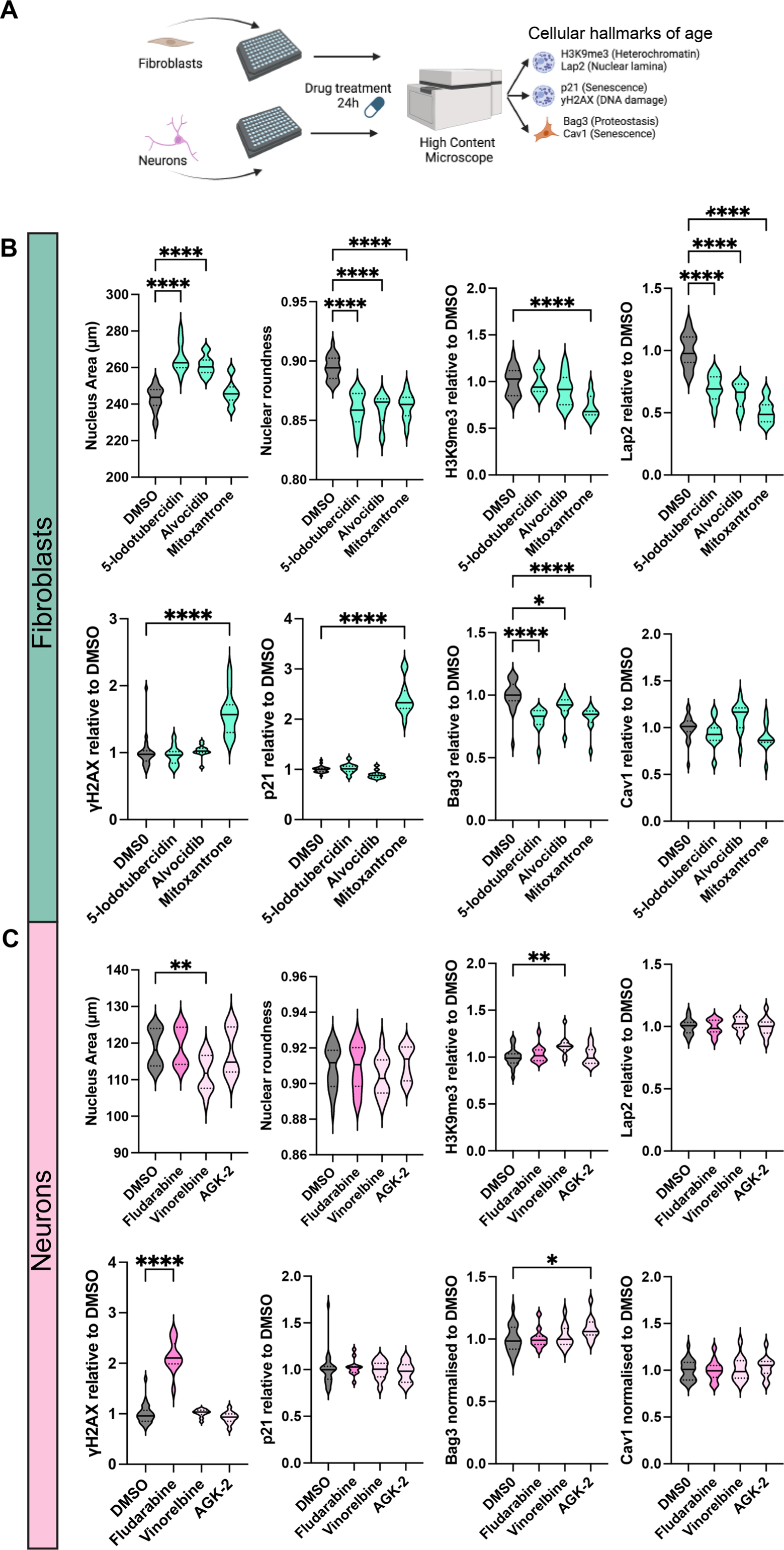
Correlation between RNAge score and cellular hallmarks of age. **(A)** Overview of experiments to measure the cellular hallmarks in neurons and fibroblasts. **(B)** Measurement of age associated cellular phenotypes in fibroblasts in response to treatment with DMSO (control), 5-Iodotubercin (10 μM), Alvocidib (10 μM) or Mitoxantrone (1.1 μM). Absolute values are shown for nuclear area and nuclear roundness. For H3K9me3, LAP2, yH2AX, p21, BAG3 and CAV1 fluorescence intensity is relative to the DMSO control. N= 12. Experiment repeated 4 times each with 3 replicate wells for both neurons and fibroblasts. **(C)** Measurement of age associated cellular phenotypes in PSC-derived neurons in response to treatment with DMSO (control), Fludarabine (10 μM), Vinorelbine (10 μM) or AGK-2 (0.04 μM). Compound with validated increase in RNAge score by RNA-seq in PSC-derived neurons (Figure 6C) are in dark pink.

Alvocidib, 5-iodotubercin and Mitoxantrone all increased the relative age of fetal fibroblasts (Figure 7B). Aging hallmarks that were induced at 24 hours post treatment across all these interventions including decreased BAG3, a loss of nuclear roundness and loss of LAP2 suggesting that decreased autophagy and nuclear lamina defects are a conserved feature of increased cellular age in fibroblasts (Figure 7B; Supp. Figure 4A). However, these were the only three assays that changed consistently across all three of our validated perturbations. Treatment with Alvocidib and 5-iodotubercin also induced an increase in nuclear area which is indicative of cellular senescence (Figure 7B). In contrast, mitoxantrone treatment increased γH2AX and p21 and decreased H3K9me3 immunoreactivity (Figure 7B). This subset of hallmarks is also strongly associated with cellular senescence. This indicates that within a specific cell type our aging score can be associated with different subsets of the aging hallmarks.

In neurons, Fludarabine had a high aging score, vinorelbine resulted in a marginal increase in the cellular age score and AGK-2 treatment had no effect as predicted by RNAge score (Figure 6C). The only aging hallmark increased in response to fludarabine treatment was the DNA damage marker γH2AX (Figure 7C and Supp. Figure 4B). In contrast, Vinorelbine treatment led to a decrease in nuclear area, which is inconsistent with cellular aging. These results suggest that neuronal aging, as determined by our transcriptional aging score, is not always associated with the complete collection of the canonical hallmarks of cellular age or that transcriptional changes precede the widespread induction of the cellular hallmarks of age.

## Discussion

Here we identify age-associated changes in transcription in the cerebral cortex, midbrain, and in primary fibroblasts and, by comparing changes in gene expression between old and young test cases, generate a tractable method for comparing cellular age between cellular conditions. This approach enables age to be scored independently of the classic cellular hallmarks of age and demonstrates that aging across different tissues and cell types involved distinct transcriptional programs and mechanisms. Drug-induced cellular age was associated with distinct transcriptional changes and aging hallmarks in neurons versus fibroblasts. Although we developed a pan-aging score and validated this score in fibroblasts, cortex, substantia nigra and cerebellum we were not able to use this score to detect an increase in the relative age of old primary stomach samples compared to young. Therefore, transcriptional aging, and perhaps the aging process more generally may be tissue specific. This idea is supported by a recent study by Buckley et al.,^8^ who developed and validated cell type specific aging clocks and showed that the different cell types in the mouse SVZ respond differently to known rejuvenation strategies. However, the transcriptomic clock developed by Buckley et al., was not suitable for our application as, due to technical limitations, they were not able to generate a clock that captured aging in neurons. Furthermore, the study was limited to the mouse and may not be directly applicable to the human CNS given the prominent species-specific differences in SVZ biology^35^. In contrast, Jung et al demonstrated that it is possible to generate a highly predictive RNAseq-based human aging clock that can be used universally across tissue types^10^. However, this method showed limited success in predicting cellular age in response to known age-related interventions, such as metformin, rapamycin, and hypoxia. This result may indicate that those interventions do not, in fact increase relative age. Alternatively, it may suggest a lack of sensitivity for a universal aging clock in detecting age-related perturbations given our evidence for cell- and tissue-specificity of aging mechanisms.

In addition to developing a novel transcriptomic measure of relative age and validating it in independent datasets, we also show its utility in tracking induced aging and rejuvenation paradigms. As human PSC-derived lineages are functionally and transcriptionally young (fetal-like) their potential in modeling late onset human disease may be limited. Age is the strongest risk factor for the several neurogenerative diseases, and there is evidence that cellular age can also impact disease progression^4,14,24,26,36^. Therefore, the ability to induce cellular aging in-vitro can provide a more physiologically relevant experimental system for early drug development and therapeutic intervention. Importantly, we show that our aging score can be used to benchmark published induced aging paradigms. For example, the ectopic expression of Progerin was able to increase relative age in the neuronal aging scores but not in the pan aging score. SATB1 loss of function in hPSC-derived midbrain dopamine neurons induced aging according to the pan, pan-neuronal and SN aging score but not the FC aging score. This is consistent with data being generated in midbrain dopamine neurons and the finding that SATB1 loss triggers cellular senescence or accelerated aging phenotypes in midbrain dopaminergic but not in cortical glutamatergic neurons. Finally, SLO treatment showed a very strong induction of neuronal age in the treated neurons triggering an increase in relative age across all the expected sub-categories.

Another intriguing finding is the maintenance of both a neuronal and fibroblast aging signature upon trans differentiation of old fibroblasts directly into neurons (iNs). Future studies will have to address to what extent a persistent fibroblast aging signature may impact the use of iNs in modeling neurological disorders and whether persistence of fibroblast age is observed across various iN induction paradigms and over prolonged iN culture periods. In contrast, we observed robust transcriptional rejuvenation signatures when comparing old primary fibroblasts with their matched iPSC-derived fibroblast counterparts, in agreement with past studies that focused on the rejuvenation of cellular hallmarks of age^3,4^. An alternative rejuvenation paradigm is the use of partial reprogramming via transient Yamanaka factor expression^18,37^. Our RNAge score should be particularly useful in tracking the dynamics of cell fate rejuvenation versus fate reprogramming under those conditions. There is broad interest in manipulating cellular age to develop novel rejuvenation strategies and increase human health span. Our transcriptomic aging score provides a platform to test and compare novel rejuvenation strategies at scale and a tool to assess therapeutic strategies for age associated diseases.

Our study demonstrates the use of aging scores to perform an in-silico screen for novel age-inducing perturbations. While our current analysis was limited to the L1000 dataset, a similar strategy could be used to probe any other pharmacological perturbation dataset with transcriptomic readouts such as Drug-Seq^38,39^ or PLATE-Seq^40^ or genetic perturbation data including possibly single cell-based strategies such as Perturb-Seq^41,42^, CRISPR-Seq^43^ or CROP-Seq^44^. The hits identified using this strategy in conjunction with the L1000 dataset had a high experimental validation rate indicating broad applicability for finding additional age-related regulators in the future. Using RNAge, the performance of novel age inducers can be directly compared using the RNAge score and tested in conjunction with previously published strategies to establish an optimal strategy to induce aging in PSC-derived neurons. Ultimately, age inducers should be combined with PSC-based disease modelling to test whether cellular age can synergize with genetic susceptibility and accelerate the onset and progression of neurodegenerative disease.

Our study demonstrates that increased transcriptional age is associated with the induction of canonical aging hallmarks in fibroblasts, whereas in neurons, the induction of those hallmarks was limited to increased DNA damage, at least upon short-term treatment. In future studies, it will be interesting to assess whether additional hallmarks of age develop in neurons following a more extended treatment regimen as this could indicate that our transcriptomic score can act as an early marker of transcriptional age. In addition, it may be important to test whether these compounds can also alter cellular age as determined by other modalities such as epigenetic aging clocks. However, this may pose challenges since DNA methylation clocks have primarily been developed using blood or saliva and are heavily impacted by cell composition rendering them difficult to use in cell culture models.

Finally, this platform provides framework to investigate the underlying mechanisms of aging across different tissues. For example, we identified fludarabine as a novel inducer of transcriptional age and showed that treating neurons with this compound resulted in increased DNA damage independent of, or prior to, the induction of any other hallmarks of age. This suggests that DNA damage may act as the primary driver of age in this specific context. In another example, we show that induction of transcriptional age in fibroblasts can be associated with two distinct patterns of cellular hallmarks of age. Mitoxantrone showed a strong cellular senescence phenotype with increased p21 expression, increased DNA damage and loss of H3K9me3 in contrast to Alvocidib and 5-Iodotubercin. This indicates that the RNAge score can capture aging triggered by distinct cellular mechanisms.

Human PSCs have become an increasingly important tool for the study of human aging and rejuvenation. Our study presents a simple strategy to score manipulations aimed at inducing, reversing or retaining cellular age across several experimental paradigms. The identification of several age-inducing and rejuvenating compounds enhances the available toolbox in the field for directing cellular age on demand.

## Acknowledgements

We would like to thank members of the Studer lab for technical support and insightful discussions. We are grateful to members of the Integrated Genomics Operation (IGO) at MSKCC and the Epigenomics Core at WCM for their excellent support in carrying out the molecular profiling assays presented in the study. The work was supported by grants from the National Institute of Aging (R01AG056298 to L.S. and R01AG054720 to L.S. and D.B, and by the core grant P30CA008748. N.S was supported in part by a Glenn Foundation for medical research postdoctoral fellowship, A.P.M. by NIA/NIH F31AG067709 fellowship and D.C. by Starr Foundation Stem Cell Scholar Postdoctoral Fellowship

## Author contributions

Conception, study design, data analysis, and interpretation, C.Z., N.S., D.C., D.B., L.S. Computational analysis and interpretation, C.Z. and D.B. Lab-based experiments were performed by N.S and D.C. with cell culture support from T.S., S.Y.C. and A.P.M. The manuscript was written by N.S., C.Z., D.B. and L.S. Funding acquired by L.S. and D.B.

## Declaration of interests

L.S. is a scientific co-founder and consultant of Bluerock Therapeutics and DaCapo Brainscience. All other authors declare no competing interests.

## Methods

### Reprogramming of primary fibroblasts to iPSCs

Young and old primary fibroblasts were obtained from Coriell and maintained in α-MEM + 15% FBS. Fibroblasts were reprogrammed to iPSCs using CytoTune Sendai viruses expressing SOX2, OCT4, KLF4 and c-MYC as previously described^4,45^. After the formation of iPSC colonies (aprox. 30 days after transduction), individual colonies were manually isolated, replated onto MEFs and maintained in KSR media.

### Differentiation of iPSCs to fibroblasts

Differentiation of iPSCs into fibroblasts was performed as previously described^4^. In brief, pluripotent stem cells were cultured on MEFs in KSR based media supplemented with 10 ng/ml FGF2. For differentiation, colonies were dissociated from the feeder layer using dispase and replated onto gelatin coated plates. Cells were maintained in DMEM + 20% heat-inactivated fibroblasts for 25 days with passaging ever 5-6 days. Fibroblasts were isolated by flow cytometry (CD-13^high^ and HLA-CD44^high^) after 25 days of culture and cultured for an additional 7 days before harvest.

### Fibroblast maintenance and culture

Fibroblasts were maintained in DMEM-F12 containing GlutaMAX,15% heat inactivated FBS and 1X Pen/Strep and passaged using Trypsin. For validation of L1000 aging score by RNA-seq old (GM04204; Coriell) or young (HDF-f; ScienCell #2300) fibroblasts were plated at a density of 4000 cm^2^, test compounds were applied when cultures reached 70% confluence.

### Generation of human cortical neurons

Cortical neurons were generated by directed differentiation of human embryonic stem cells (H9; WA-09) maintained in E8 on vitronectin coated plates^46^. Differentiation was performed by seeding pluripotent stem cells onto Matrigel coated dishes at a density of 300,000/cm2 in the presence of Y27632 (10□μM). After 24h the E8 medium was replaced with E6 containing SB431542 (10□μM), LDN193189 (100□nM) and XAV939 (2□μM). After three days the XAV939 was removed from the differentiation media and cells were cultured for an additional seven days in E6 containing SB431542 (10□μM), LDN193189 (100□nM). The neuroepithelium was maintained for an additional 10 days in neurobasal supplemented with N2 and B27. Media was changed daily throughout the differentiation. At DIV 20 the cultures were dissociated using Accutase to generate a single cell suspension and plated out for subsequent experiments in Neurobasal medium supplemented with B27, L-glutamine, BDNF, cAMP, ascorbic acid and GDNF. DAPT was also added to the culture medium from DIV20 to DIV30.

### Compounds for experimental validation of L1000-based score

All validation experiments were performed after 24h of compound treatment. For rejuvenation experiments in fibroblasts Mocetinostat (10 μM; Selleck S1396), Resveratrol (0.37 μM; Selleck S1396) and Radicicol (1.1 μM; Tocris 1589), DMSO (1:1000) and EtOH (1:1000; vehicle control for Radicicol) were applied to old primary fibroblasts (Coriell GM4204). Age induction experiments in young fibroblasts (HDF-f; ScienCell #2300) were performed with 5-Iodotubercin (10 μM; Selleck S8314), Alvocidib (10 μM; Selleck S1230) and Mitoxantrone (1.1 μM; Selleck S2485) with DMSO (1:1000) used as a control. Compounds used for age induction experiments in PSC-derived neurons were: Fludarabine (10 μM; Selleck S1491) Vinorelbine (10 μM; sc-205885) and AGK-2 (0.04 μM; Selleck S7577) with DMSO (1:1000) used as a control.

### Assaying Hallmarks of Age by high content microscopy

HDF fibroblasts were plated at a density of 4500/cm^2^ and cultured for 48h before drug treatment. Neurons were plated at a density of 100,000/cm^2^ and cultured for 12 days before the application of candidate age inducers. After 24h of treatment culture wells were washed 2x in PBS then fixed with 4% paraformaldehyde. For immunocytochemistry cells were permeabilized in PBS+0.3% Triton and blocked in 5% normal goat serum. All antibody primary incubations were performed overnight at 4 degrees and secondary antibodies at RT for 1h. Primary antibodies used in this study were: BAG3 (abcam; ab47124), CAV1 (Thermo; MA3-600), H3K9me3 (abcam; ab176916), LAP2 (BD Bioscience; 611000), p21 (CST; 2947), yH2AX (EMD Millipore; 05-636). Immunocytochemistry images were acquired using an Operetta (PerkinElmer, Waltham, MA) microscope and quantified using the harmony high content analysis software. For both fibroblasts and neurons 4 independent batches of cells were assayed in triplicate (n=12). Fluorescence intensity was normalized to DMSO, and ordinary one-way ANOVA used to compare test conditions to the DMSO control (Prism 9.5.1).

### Flow Cytometry

Cell sorting for iPSC derived fibroblasts was performed using CD13-PE and CD44-APC conjugated primary antibodies. Antibodies were incubated with 1×10^6^ cells on ice for 1h and sorting was performed using FACS Aria (BD Biosciences).

### RNA Extraction

Cell were harvested, lysed, and stored in Trizol (Life Technologies) until further processing. For primary frontal cortex or substantia nigra a total of 10 mg of brain tissue from was lysed in Trizol aided by a tissue dounce. For experimental validation L1000 compounds RNA was extracted by the core using miRneasy Micro Kit (QIAGEN 1071023). In all other cases RNA was extracted using the Zymo Direct-zol RNA Microprep Kit according to the manufacturer’s instructions.

### DNA Extraction

DNA was extracted using the DNeasy Blood & Tissue Kit (Qiagen), with RNAse step performed for 10mins. Eluted DNA was concentrated using Genomic DNA clean and concentrator (Zymo).

### RNA-seq analysis

Reads were aligned to the hg19 Human transcripts using STAR^47^ (version 2.5.21b) using default parameters and resulting bam files were sorted and indexed using samtools. Gene counts were obtained using featureCounts^48^ (version 1.4.3) from sorted bam files using uniquely mapped reads. Genes with no expression counts in any sample were discarded. Differential gene expression analysis was performed using DESeq2^49^ R package that normalize gene count data to transcription per million (TPM), and then detect differentially expressed genes (DEG) between Young and Old groups with (FDR < 0.1).

### 5hmc data analysis

Reads were aligned to the hg19 Human genome using Bowtie2 (V2.2.5)^50^ with default parameters and resulting BAM files sorted and index using samtools. MACS (V 2.1.0)^51^ with default parameters used to detect statistically significant peaks by identifying genomic regions with enriched coverage in ChIP samples relative to input control.

### 5mc data analysis

FASTQ files were generated by bcl2fastq (V2.17) and filtered for pass filter reads based on Illumina’s chastity filter. Sequencing adapters were trimmed by FLEXBAR (V2.4)^52^, genomic alignments using Bismark (V0.14.4)^53^ and Bowtie2 (V2.2.5)^50^ to reference human genome hg19, and per base CpG methylation metrics were calculated with a custom PERL script. CpGs at a minimum threshold coverage of 5 reads were used for downstream analysis.

### Functional enrichment analysis

Mosaic version 1.1 was used to retrieve gene ontology (GO) information for all genes of the Human genome^54^. Functional analysis was performed on DEG with DAVID^55^ (version 6.8) and biological process GO terms and KEGG pathways with enrichment p < 0.05 were selected as overrepresented functions. To remove the redundancy of GO terms, up to top 350 GO terms were used as the input of REVIGO (http://revigo.irb.hr/) to generate the list of intelligible functions.

### RNAge Training datasets

Total pF: Primary fibroblast samples from 9 young (7-14 years old) and 9 old (70-96 years old) profiled by total-RNA-seq protocol.

PolyA pF: Randomly picked 4 young (10-11 years old) and 4 old (71-96 years old) out of above 18 primary fibroblast samples. These samples were profiled by polyA RNA-seq kit.

PolyA Gage pF: Primary fibroblast RNA-seq data from 6 young (< 15-year-old) and 4 old (> 70-year-old) were downloaded from E-MTAB-3037 dataset^13^. All raw data were processed by our RNA-seq analysis protocol.

PolyA Gage FC: frontal cortex RNA-seq data from 4 young (< 15-year-old) and 4 old (> 70-year-old) were downloaded from E-MTAB-3037 dataset^13^. All raw data were processed by our RNA-seq analysis protocol.

Total FC: Primary frontal cortex samples from 9 young (13-14 years old) and 9 old (70-91 years old) profiled by total-RNA-seq protocol.

Total SN: Primary substantia nigra samples from 10 young (13-14 years old) and 9 old (70-91 years old) profiled by total-RNA-seq protocol.

### Other datasets

GSE113957: Fleischer et al. collected fibroblast samples from 133 healthy individuals^9^. The FPKM of gene expression were downloaded from GEO database. Out of 133 samples, 10 young (8-13 years old), 10 old (> 89-year-old) and 10 HGPS (2–8 years old) samples were used in this study.

GSE36192: Microarray data of the cerebellum and frontal cortex from 396 subjects (total 911 tissue samples) were generated by North American Brain Expression Consortium^16^. The normalized gene expression matrix was downloaded from GEO database. Both cerebellum and frontal cortex from the same 18 young (10-15 years old) and 16 old (> 92-year-old) samples were used in this study.

GSE52431: The raw RNA-seq data from 4 iPSC-derived dopamine neurons with overexpressing progerin and 4 corresponding control samples were downloaded and processed with proposed pipeline^4^.

GSE132040: Bulk RNA-seq data from Tabula Muris Consortium study were downloaded and processed. The whole brain data from young (<3-month) and old (>24-month) mice were used in this study^56^.

GSE141028: Fathi et al. presented a study of inducing senescence by treating human embryonic stem cells with compounds^26^. The raw RNA-seq data from 3 treatment and 3 WT cell lines were downloaded, processed, and used in this study.

E-MTAB-5965: Genetic ablation of SATB1 induces a senescence phenotype in human embryonic stem cell (hESC)-derived DA neurons^25^. The raw RNA-seq data from 3 STAB1-KO and 3 WT cell lines were downloaded, processed, and used in this study.

E-MTAB-10352: The raw RNA-seq data from 42 samples were downloaded, processed, and used in this study^14^, including 17 samples from iPSC derived induced neurons and 25 samples from direct conversion of human fibroblasts into induced neurons.

GTEx: The FPKM data of four tissues was downloaded from GTEx Portal^17^, including frontal cortex, cortex, substantia nigra and stomach.

### Aging markers identification

We derived the aging signature for the different tissue types using the following six datasets:

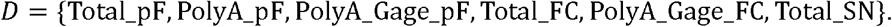

Differential gene expression analysis was performed using DESeq2 between young and old samples in each of the RNA-seq datasets, resulting in log fold change values and p-value from each of the six datasets.

- The log2 based fold change between young and old groups for each gene *i* in each dataset *j* is denoted as *L*_*i,j*_, where *i* ∈ *G,j ∈ D*.
- The p-value between young and old groups for each gene in each dataset is denoted as *p*_*i,j*_, where *i* ∈ *G,j ∈ D*.

We discretized the log2 fold change of the genes to indicate the regulatory direction in aging. We defined a simplified log2 fold change as follow:

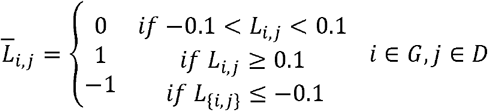

Then we combined p-values from all six dataset for each gene with Edgington’s method ^57^:

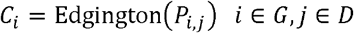

Based on the simplified 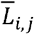, we assign genes to discrete and mutually exclusive aging signatures in hierarchical scheme as follow:

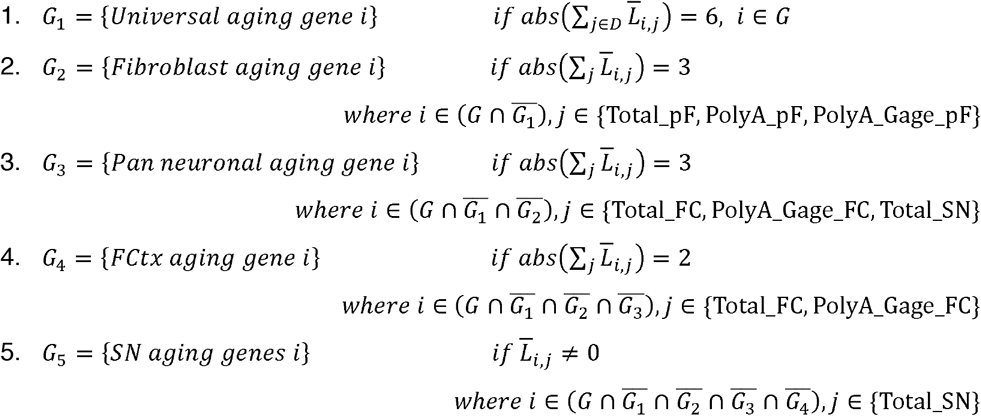

For each aging signature set, we ranked genes based on their combined p-value *C*_*i*_ . Thus top 100 genes with smallest p-values from each subset were selected as the markers for the downstream analysis. These top ranked gene subsets denoted as 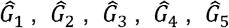, respectively. According to the definition, the simplified 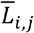 of above signatures is consistent across multiple datasets from the corresponding tissue type and it excludes genes that do not exhibit any change between the young and old groups. We can further define a marker vector for a tissue type *k* as

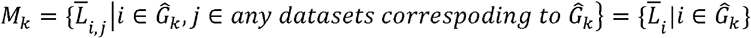

### RNAge score calculation

The aging score is applied on a gene expression test data where two conditions are compared for expression changes in the aging signature, group1 has *n* samples and group2 has *m* samples. For each gene in a tissue specific marker set 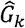, we can calculate t-statistic between two groups based on Welch’s two-sample t test as follow:

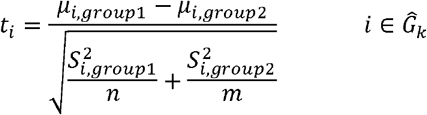

where *μ*_*i,group*1_ is the mean expression of gene *i* in group1, *μ*_*i,group*2_ is the mean expression of gene *i* in group2, 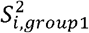 is the variance of gene *i* in group1, and 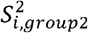 is the variance of gene *i* in group2.

All t-statistics from the same tissue specific marker set 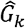 denoted as a vector 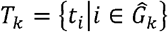 In cases where marker gene *i* is not expressed in the test dataset we set *t*_*i*_=0. Then we defined the “Aging score” for given tissue type *k* as follow:

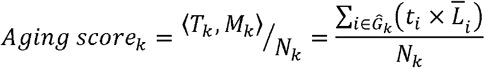

where the normalizing factor *N*_*k*_ is the number of overlapping genes between the test dataset and 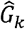. The aging score, *Aging score*_*k*_, represents the overall magnitude of expression difference between two sample groups based on the expression trends. This score is calculated using the tissue-specific aging marker set *M*_*k*_.It indicates the aging difference between the two groups as well as distinguishes which group is older. To mitigate the influence of a small number of markers with large changes gene expression between group1 and group2 of the test data (i.e. large t-score values), we introduced the “Percentage score” as a measure of agreement between the expression trends of the aging markers and the given dataset. The calculation of the Percentage score involves the product of each gene between and *T*_*k*_ and *M*_*k*_: 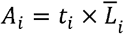, where 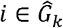. A positive *A*_*i*_ indicates a consistent expression change of gene *i* in both the aging signature and in the comparison between group1 and group2 in the test dataset. Conversely, a negative *A*_*i*_ indicates an opposite expression trend. Therefore, the “Percentage score” is defined as:

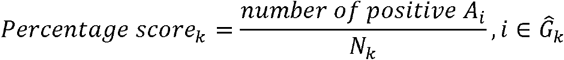

### Data and software availability

RNA-seq, 5mc and 5hmc data used in this paper are available at NCBI Sequence Read Archive (https://www.ncbi.nlm.nih.gov/sra) under accession SRA: PRJNA1002866 and PRJNA1003556. Source code files for generating figures are available online (https://github.com/zhangch/RNAgePaper).

## Notes

### Summary of Updates

Author information updated; Minor changes to text and figures.

