## Supplemental Figures for "Identifying novel age-modulating compounds and quantifying cellular aging using novel computational framework for evaluating transcriptional age"

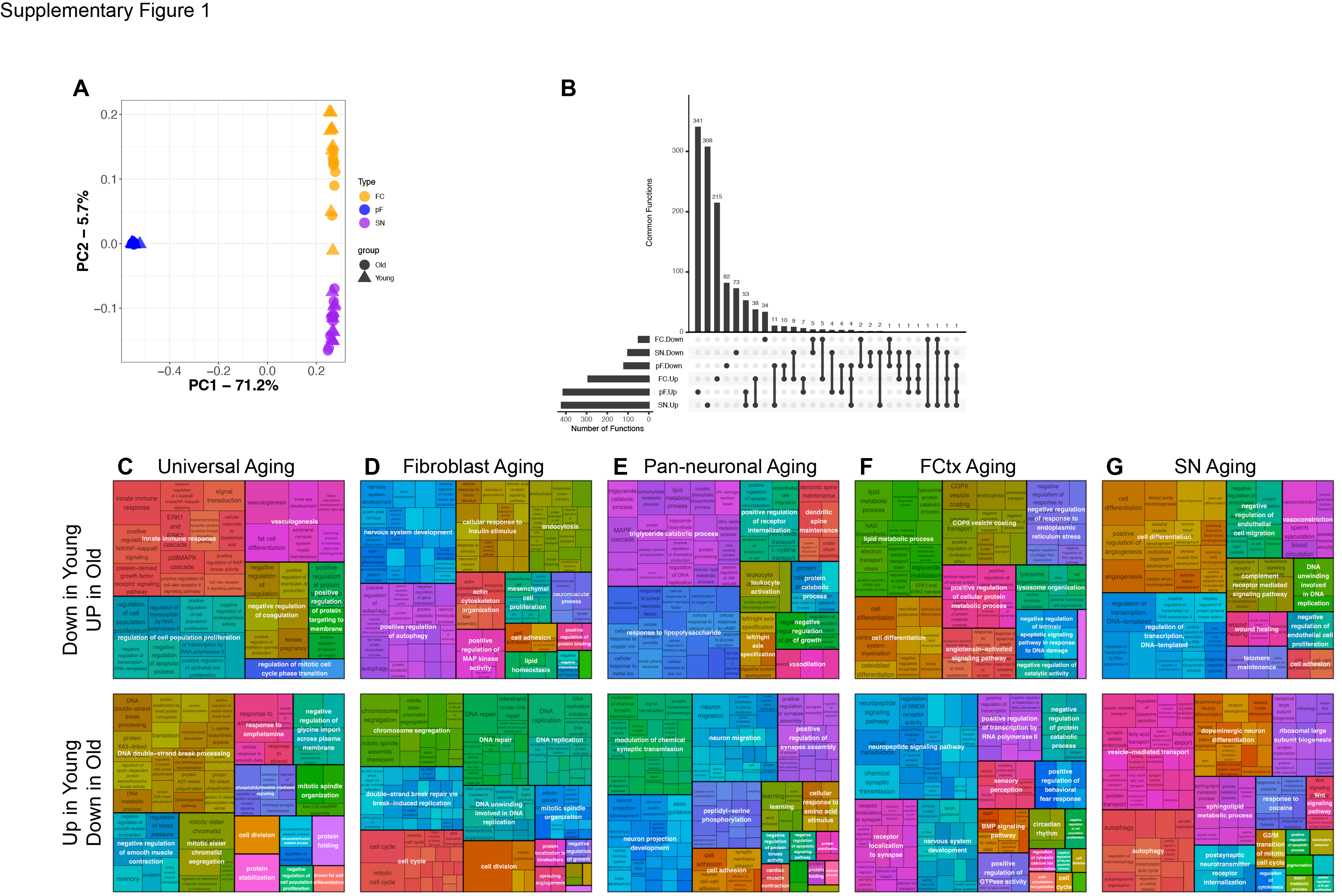

**Supp. Figure 1: Functions associated with aging in primary fibroblasts, FC and SN**

**(A)** PCA plot of all samples sequenced for this experiment. FC= frontal cortex, pF = primary fibroblast, SN= substantia nigra. **(B)** Plot showing the number of overlapping functions associated with aging in FC (frontal cortex), SN (substantia nigra) and pF (fibroblasts). Down indicates that genes associated with this function are expressed at lower levels in young than old. Up indicates that genes associated with this function are expressed at higher levels in young than in old. **(C-G)** Functional enrichment of genes associated with each of our aging signatures from Figure 1E.

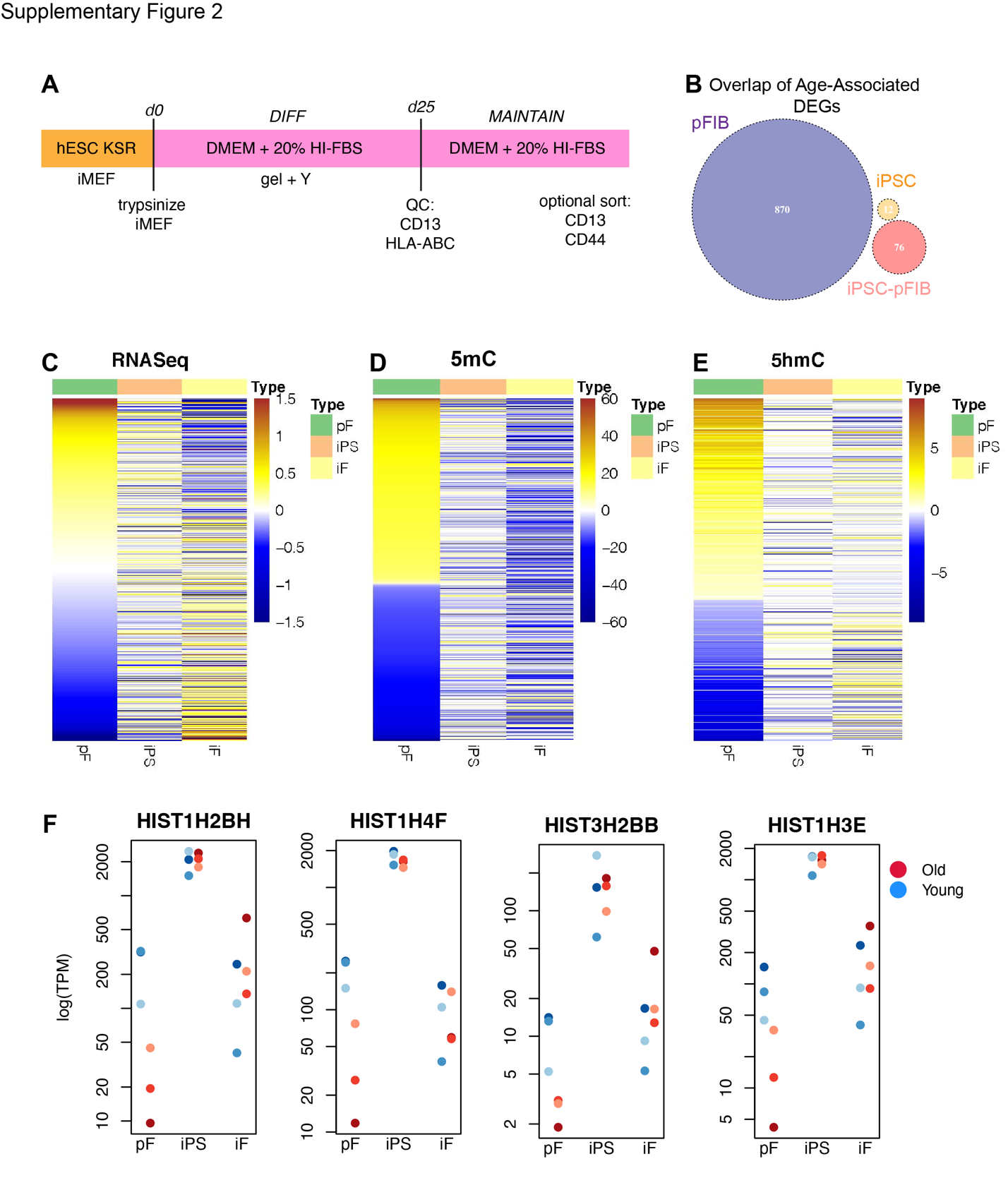

**Supp. Figure 2: Extended characterization and validation of iPSC reprograming as a model of cellular rejuvenation.**

**(A)** Schema outlining the protocol used to generate iPSC-FIBs. **(B)** Venn diagram of DEGs between iPSCs and iPSC-FIBs of young and old origin indicates lack of overlap between the DEGs in the pFIBs.
Heatmaps showing (**C**) differential gene expression, (**D**) differentially methylated promoters (5mC) and (**E**) differences in hydroxymethylation (5hmC) between young and old pFIBs (N=4 independent cell lines derived from young donors and 4 derived from old donors), iPSCs of young and old origin N=3 independent cell lines derived from young donors and 3 derived from old donors) and matched iPSC-FIBs (N=3 independent cell lines derived from young donors and 3 derived from old donors). **(F)** Normalized expression of histone genes (*HIST1H2BH*, *HIST1H4F*, *HIST3H2BB* and *HIST1H3E*) in pFIBs (pF), iPSC-FIBs (iPS) and iPSC-FIBs (iF).

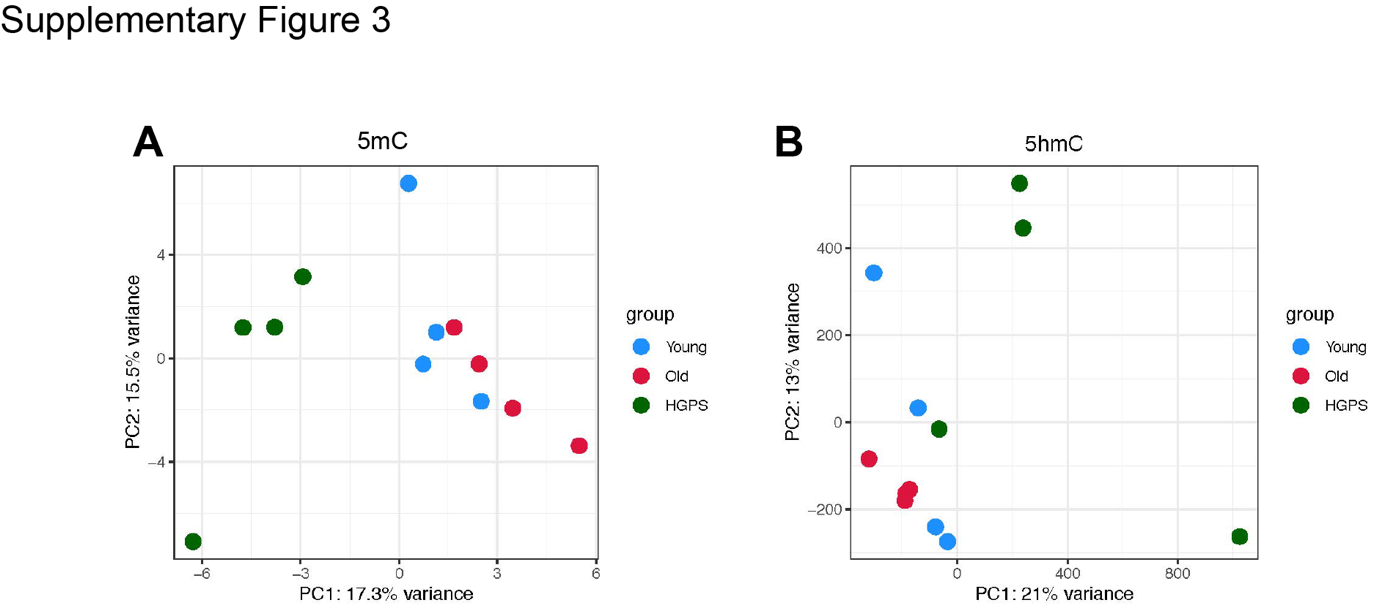

**Supp. Fig 3: DNA methylation analyses of young, old and HGPS fibroblasts.**

**(A)** PCA of promotor 5mC data from primary fibroblasts originating from young, old or individuals with HGPS. **(B)** PCA of 5hmC data from primary fibroblasts originating from young, old or individuals with HGPS.

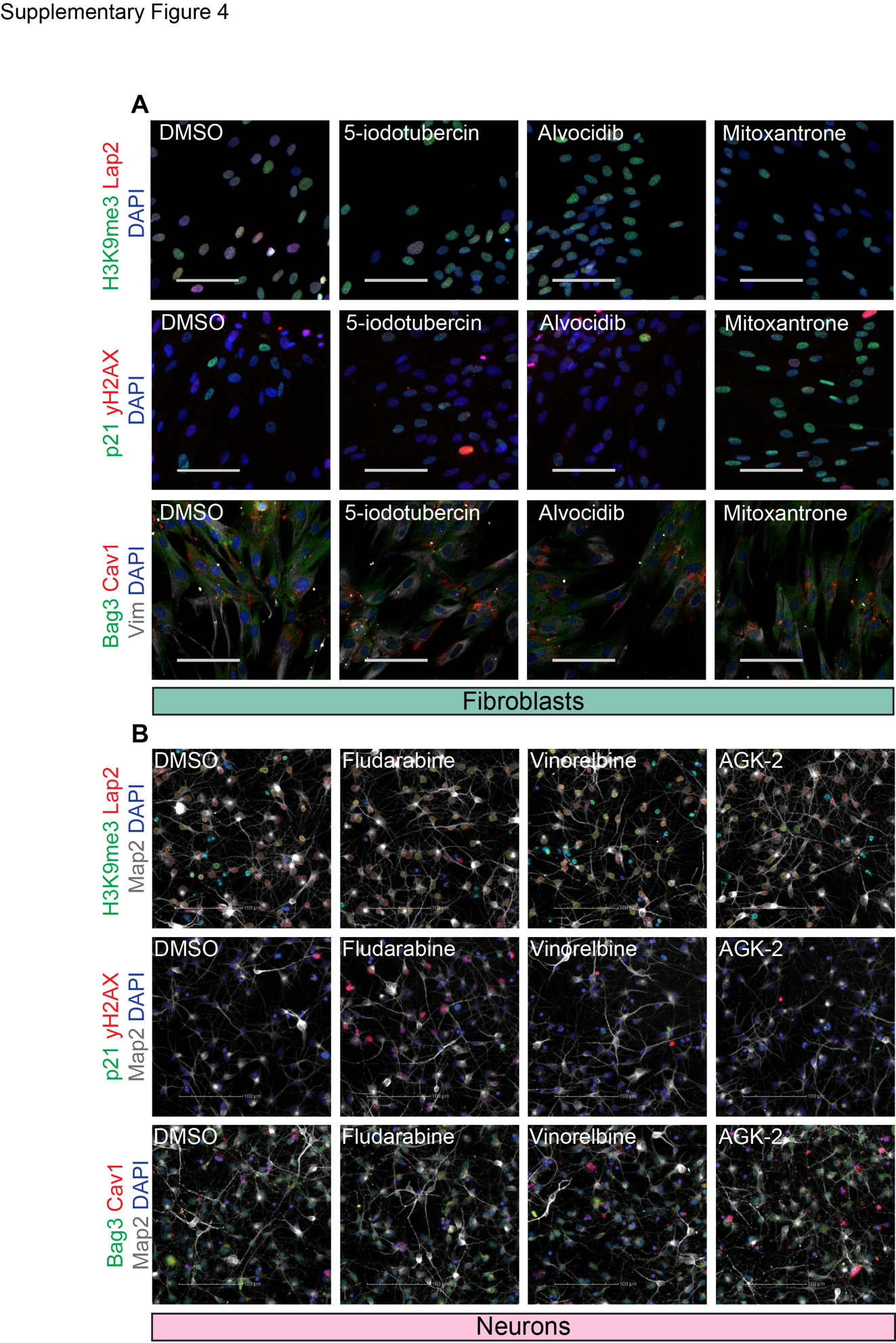

**Supp. Figure 4: Example immunocytochemistry images for the hallmarks of age assays performed in fibroblasts and PSC-derived neurons.**

**(A)** Example immunocytochemistry images of hallmarks of aging assays in fibroblasts. **(B)** Example immunocytochemistry images in hallmarks of aging assays performed in PSC-derived neurons. Scale 100uM
